# Foundational characterization of tomato fruit RALF peptides reveals structural and functional specialization within the SlRALF family

**DOI:** 10.64898/2026.06.05.730363

**Authors:** José A. Montano, Marta Carrera, Xu Wang, Pablo Mesa-Rojas, Ana M. Luna, Andreas Schaller, Ian Morilla, Verónica G. Doblas

**Affiliations:** Instituto de Hortofruticultura Subtropical y Mediterránea La Mayora, Universidad de Málaga-Consejo Superior de Investigaciones Científicas (IHSM, UMA-CSIC), Málaga, Spain; Department of Plant Physiology and Biochemistry, Institute of Biology, University of Hohenheim, Stuttgart, Germany

## Abstract

Rapid Alkalinization Factor (RALF) peptides regulate plant growth and cell wall signaling, but their roles in fruit development remain unclear. Here, we characterized tomato (*Solanum lycopersicum*) fruit-associated RALF peptides and their interactions with leucine-rich repeat extensins (LRXs). Expression analyses identified *SlRALF5*, *SlRALF7*, and *SlRALF10* as the main fruit-expressed RALFs. *SlRALF10* was associated with early fruit development, whereas *SlRALF5* and *SlRALF7* remained expressed during ripening. Sequence analyses showed that SlRALF5/7 retain conserved motifs of canonical RALFs, while SlRALF10 displays divergent structural features and altered charge distribution. Synthetic SlRALF5 and SlRALF7 inhibited root growth and induced extracellular alkalinization, whereas SlRALF10 lacked both activities. Co-immunoprecipitation assays showed that all three peptides interact with the fruit-expressed proteins SlLRX2 and SlLRX5. Structural modeling predicted distinct electrostatic properties for the SlLRX5/SlRALF10 complex compared with SlRALF5. These results reveal structural and functional specialization among tomato fruit RALF peptides and suggest that distinct SlRALFs may differentially respond to cell wall remodeling during fruit development and ripening.

## Introduction

RAPID ALKALINIZATION FACTOR (RALF) peptides constitute a family of cysteine-rich plant peptide hormones that are involved in a wide range of biological processes, including root growth, reproduction, immune responses, and adaptation to abiotic stress conditions (Campbell & Turner, 2017; Lu et al., 2025; Murphy & De Smet, 2014; Pratyusha & Sarada, 2025; Schade et al., 2025). RALF peptide precursors typically range from 50 to 250 amino acids in length and generally contain an N-terminal signal peptide that directs them into the secretory pathway, where they undergo proteolytic maturation mediated by Site-1 Protease (S1P), a subtilisin-like serine protease that recognizes a RRXL motif located upstream of the RALF active domain (Abarca et al., 2021; Blackburn et al., 2020; Pearce et al., 2010; Srivastava et al., 2009). RALF peptides have been identified in more than 50 plant species, underscoring their evolutionary conservation and functional relevance (Campbell & Turner, 2017; Murphy & De Smet, 2014; Zhang et al., 2020; 2023).

RALF peptides were initially identified as ligands for plasma membrane-localized *Catharanthus roseus* receptor-like kinases 1-like (CrRLK1Ls) proteins and glycosylphosphatidylinositol (GPI)-anchored proteins of the LORELEI/LORELEI-like GPI-anchored protein (LRE/LLG) family (Solis-Miranda & Quinto, 2021; Zhu et al., 2021). More recently, RALF peptides have been shown to play dual structural and signaling roles in maintenance of cell wall integrity (Manchanda & Geitmann, 2025). RALFs interact with cell wall-associated receptors known as Leucine-rich repeat (LRR)-extensin (LRX) proteins. These peptides exhibit high affinity for the LRR domain under acidic conditions, whereas the extensin domain is covalently cross-linked to the wall matrix (Herger et al., 2019; Moussu & Ingram, 2023). The C-terminal region of RALFs binds to LRX proteins, exposing positively charged cationic residues that physically interact with de-methylesterified, negatively charged pectins, thereby forming supramolecular assemblies that contribute to cell wall integrity during plant development (Moussu et al., 2023; Schoenaers et al., 2024). In Arabidopsis, the RALF4-LRX8 complex binds specifically to de-methylesterified homogalacturonan (HG) in a charge-dependent manner, thereby influencing cell wall organization during pollen tube growth (Moussu et al., 2023). Similarly, the RALF22-LRX1/2 complex organizes de-methylesterified HG into circumferential rings at the root hair tip, modulating cell wall architecture (Schoenaers et al., 2024).

De-methylesterified pectins promote cell wall loosening and increase cell wall permeability, processes associated with cell expansion and elongation in structures such as root hairs and pollen tubes. Given that fruit ripening also involves extensive de-methylesterification and remodeling of HG, similar RALF-mediated mechanisms may operate in fruit tissues. Pectins are particularly abundant in fruit cell walls, accounting for up to 60% of the cell wall dry weight, and their remodeling is a key feature associated with fruit softening (Brummell, 2006; Hyodo et al., 2013; Posé et al., 2019; Shi et al., 2023). Fruit softening is generally characterized by highly coordinated pectin remodeling. HG is secreted into the apoplast as long chains in a highly methyl-esterified state. Pectin methylesterases (PMEs) are typically upregulated during ripening; for example, in tomato, pectins are approximately 90% methyl-esterified in mature green fruit, decreasing to around 35% in red-ripe fruit (Hyodo et al., 2013; Jeong et al., 2018). PMEs can act in a random manner, rendering HG more susceptible to degradation by pectinolytic enzymes such as polygalacturonase (PG) and pectate lyase (PL) (Palin & Geitmann, 2012). Consistently, downregulation of PG or PL genes has been shown to enhance shelf life and/or reduce fruit softening in tomato, strawberry, apple and peach (Atkinson et al., 2012; Paniagua et al., 2016; Qian et al., 2021; Santiago-Doménech et al., 2008; Uluisik & Seymour, 2020; Wang et al., 2018). Understanding the regulatory mechanisms underlying pectin remodeling is therefore of major agronomic interest for improving fruit firmness and postharvest shelf life.

The involvement of RALF peptides in pectin remodeling during fruit development has received limited attention (Huang et al., 2024; Montano et al., 2026). One of the few studies available was conducted in strawberry, where silencing the expression of *FaRALF29* delayed fruit colonization by *Colletotrichum* species, pathogens associated with fruit rot during storage and transport (Cesarino, 2023). In contrast, overexpression of the *FaRALF29* gene in unripe white fruits induced disease symptoms (Merino et al., 2019). We hypothesized that fruit-expressed RALF peptides regulate pectin remodelling and contribute to fruit softening during tomato ripening. Transcriptomic analyses identified *SlRALF5*, *SlRALF7* and *SlRALF10* as preferentially expressed during fruit development. In this study, as a first step towards understanding their role, we characterized them through physiological and biochemical approaches.

## Results

### *SlRALF* and *SlLRX* gene expression pattern analysis

The tissue-specific expression of tomato *CrRLK1L* genes has been reported by Ma et al., 2023. Among the 24 *SlCrRLK1L* genes, three showed high expression in fruit tissues: *SlCrRLK1L20* was the most highly expressed, reaching its maximum at the breaker (Br) plus a 10-day fruit stage; *SlCrRLK1L15* exhibited a similar pattern to *CrRLK1L20*, but with lower expression levels; and *CrRLK1L7* showed a moderate expression in fruit tissue, peaking at the Br stage and decreasing at Br+10 (Ma et al., 2023). To investigate the tissue-specific expression of tomato *SlRALF* and *SlLRX* genes, we followed a similar approach. The expression profiles of all the 11 *SlRALF* genes and 5 *SlLRX* genes were analyzed in the following tissues: root, leaf, bud, flower, 1-3 cm fruit, mature green fruit, breaker fruit, and breaker plus a 10-day fruit. The original RNA-seq data were obtained from the SGN tomato functional genomic database (SGN-TFGD, http://ted.bti.cornell.edu/cgi-bin/TFGD/digital/home.cgi) (Sato et al., 2012). As shown in Figure 1a, *SlRALF5* and *SlRALF7* genes were predominantly expressed at the mature fruit stage. *SlRALF9* and *SlRALF10* were expressed up to mature green stage, after which their expression declined. The remaining *SlRALF* genes showed little or no expression in fruit tissue, appearing to be specific to other organs, for example *SlRALF2* in roots, and *SlRALF1*, *SlRALF3*, *SlRALF6*, *SlRALF8* in buds and flowers. As shown in Figure 1b, two *SlLRX* genes were expressed in fruit tissue: *SlLRX5* was the most highly expressed, peaking toward the end of ripening, while *SlLRX2* exhibited lower expression levels. *SlLRX1* and *SlLRX3*, in contrast, were mainly expressed in flowers. Based on these expression pattern, we hypothesize that *SlRALF5* and *SlRALF7* represent candidate regulators during the ripening process, whereas *SlRALF9* and *SlRALF10* may function during cell expansion phase up to the mature green stage.

**Figure 1:**
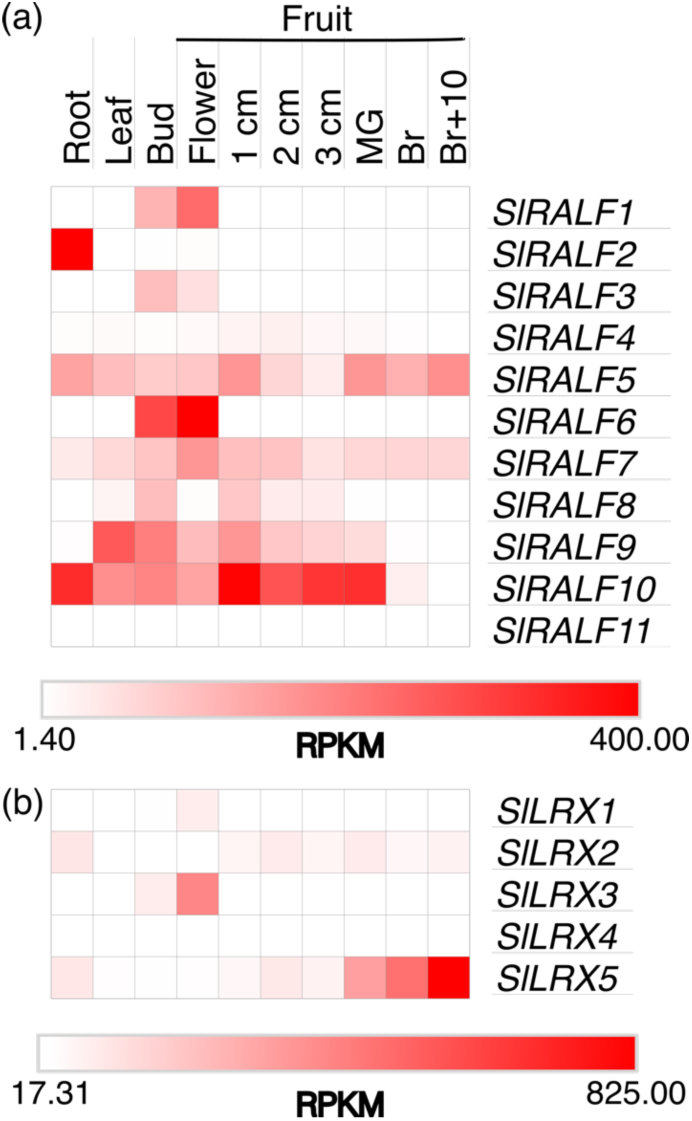
Expression pattern analysis of *SlRALFs* and *SlLRXs*. (a) Heatmap of *SlRALF* expressions in a variety of tissues. (b) Heatmap of *SlLRX* expressions in a variety of tissues. RPKM: reads per kilobase per million mapped reads. All of the data were acquired from the SGN RNA-seq database.

*Sl*RALF5 and *Sl*RALF7 cluster within the same group as Arabidopsis RALF1, RALF22, RALF23 and RALF33, which also includes most fruit-expressed RALFs, such as *Fa*RALF1 and *Fa*RALF39, both of which show the highest expression levels in mature strawberry fruits (Montano et al., 2026; Negrini et al., 2020). In contrast, *Sl*RALF9 and *Sl*RALF10 clusters within a less well-supported group together with Arabidopsis RALF27 (Montano et al., 2026). Notably, *Sl*RALF10 was not included in the classification proposed by (Campbell & Turner, 2017). To investigate the relationship between Arabidopsis and tomato LRX proteins, a phylogenetic analysis was performed using full-length amino acid sequences (Supplementary Figure 1). *Sl*LRX5, which exhibits the highest expression during fruit ripening (Figure 1b), groups together with Arabidopsis LRX5, expressed in roots and shoots, and LRX6, which is associated with lateral root formation (Supplementary Figure 1) (Herger et al., 2019). *Sl*LRX2, which also shows moderate expression during fruit ripening, cluster with Arabidopsis LRX1 and LRX2, both predominantly expressed in root hairs (Supplementary Figure 1) (Herger et al., 2019).

### Conserved motifs in the mature forms of *Sl*RALF5, *Sl*RALF7 and *Sl*RALF10 proteins expressed in tomato fruit

To investigate the role of RALFs in fruit formation, we selected *SlRALF5* and *SlRALF7* as candidates of interest, as they are expressed during fruit ripening, and *SlRALF10*, which shows high expression from the initial stages of fruit development until the mature green stage (Figure 1a). Phylogenetic analysis revealed that SlRALF5 and SlRALF7 proteins share high sequence similarity with SlRALF2, which is expressed in roots (Montano et al., 2026) (Figure 1a). These three *Sl*RALF peptides group together with Arabidopsis RALF1, RALF22, RALF23 and RALF33 (Montano et al., 2026). We have also included Arabidopsis RALF4 and RALF19, since they have been well characterized with respect to amino acid motifs important for their function (Moussu et al., 2020; 2023; Schoenaers et al., 2024; Xiao et al., 2019). SlRALF9 and SlRALF10 cluster with Arabidopsis RALF27 (Montano et al., 2026). We aligned all mature form from protein sequences to compare them with the motifs described in the literature (Figure 2).

**Figure 2:**
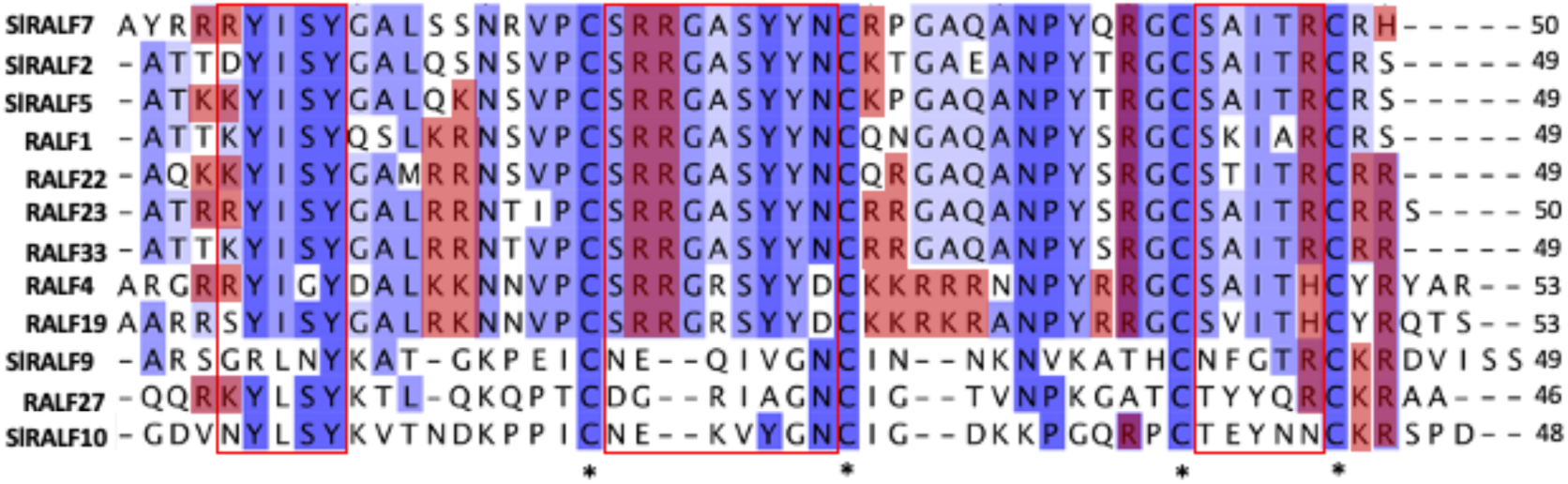
Alignment of the mature sequences of SlRALF5, SlRALF7 and SlRALF10 with their related tomato and Arabidopsis RALF members. Conserved residues are highlighted in different blue tones, whereas basic residues are shown in red. Motifs reported to mediate interactions with LLG proteins (N-terminal region) and LRX proteins (middle and C-terminal regions) proteins are enclosed in red boxes. Conserved Cys residues are indicated by black stars.

SlRALF2, SlRALF5 and SlRALF7 are classified as true RALFs, as their sequences contain the RRXL cleavage site, the YISY motif, and the other well conserved motifs GASYY, PYXRGCS and RCRR(S) (Figure 2) (Bedinger et al., 2010; Campbell & Turner, 2017). The YISY motif is essential for RALF1-induced alkalinization, together with the Leu-11 residue, whose deletion was reported to have a profound effect on alkalinization activity (Pearce et al., 2010). SlRALF2, SlRALF5 and SlRALF7 conserve this Leu residue, whereas SlRALF9 and SlRALF10 do not (Figure 2). Residues 4-17 at the N-terminal of RALF23 establish extensive interactions with LLG2, where only the YISY motif is required to nucleate the ternary complex LLG-RALF-FER (Xiao et al., 2019). The N-terminal residues of RALF23 reported to form polar interactions with LLG2 are conserved in SlRALF2, SlRALF5 and SlRALF7 (Figure 2). However, the basic Arg residues surrounding the YISY motif in RALF23, important for polar interaction with LLG2, are not fully conserved. SlRALF2 lacks these residues entirely, while in SlRALF5 and SlRALF7 some are replaced by polar amino acids (Figure 2). Among the other Arabidopsis RALFs (RALF1, RALF4, RALF19, RALF22 and RALF33), these residues are largely conserved, with only minor substitutions in some cases. SlRALF10 possesses an equivalent YLSY motif, similar to Arabidopsis RALF27, whereas SlRALF9 lacks the YISY/YLSY motif entirely (Figure 2). The basic Arg residues surrounding the YISY/YLSY motif are also absent in SlRALF10, making it more similar to SlRALF2.

Analysis of the RALF4-LRX8 interface revealed that LRX proteins recognize the mature sequence, and the YISY motif is shown to be unnecessary for binding (Moussu et al., 2020). All four cysteines in RALF4 were required for proper peptide folding and high affinity binding to LRX8 (Moussu et al., 2020). These cysteines are conserved across all five SlRALF peptides (Figure 2). Two loops formed by the cysteine pairs are critical for LRX8 recognition (Moussu et al., 2020). The first loop is highly conserved among SlRALF2, SlRALF5 and SlRALF7 and related Arabidopsis RALFs. It contains two tyrosine residues that directly contact the LRR core (Figure 2) (Moussu et al., 2020). These tyrosines are conserved in all analyzed RALFs except RALF27 and SlRALF9, where nonpolar residues occur, and in SlRALF10, where one tyrosine is replaced by a Glycine (Figure 2). Apart from a few charged residues specific to RALF4 and RALF19, this loop is otherwise conserved. The second shows a similar pattern, with a single charge residue unique to RALF4 and RALF19 (Moussu et al., 2020). Both loops are absent in SlRALF9 and SlRALF10 (Figure 2).

Between the two cysteines loops, RALF4 exhibits an exposed, positively charged surface when bound to LRX8, composed by residues Arg32-34 (89-91) in a protruding loop and extending to Lys13 (70), Arg21 (78), Lys30-31 (87-88) and Arg39 (96). Numbers in brackets are from the N-terminal part of the full prepropeptide according to (Moussu et al., 2020; 2023). This basic patch mediates electrostatic interaction with pectin (Moussu et al., 2020; 2023). Positively charged residues at positions 4-5 (61-62), located immediately upstream of the YISY motif, are conserved in RALF22, RALF23, RALF19, RALF27, SlRALF5 and SlRALF7, but absent in SlRALF9 and SlRALF10 (Figure 2). Lys13 (70) is not conserved in SlRALF2, SlRALF5 or SlRALF7, whereas Arg21 (78) is fully conserved (Figure 2). The main positively charged patch in RALF4 (residues 30-34) is fully conserved only in RALF19; in other RALFs, these residues are replaced by polar and nonpolar amino acids (Figure 2). While RALF22 retains only one Arg of this positively charged patch, most of the other basic residues are conserved with RALF4, including three C-terminal Arg residues, but within positions 30-34 (87-91) only one Arg remains (Schoenaers et al., 2024). SlRALF5 and SlRALF7 show a pattern similar to RALF22, except that Arg82 and Arg100 are not conserved (Figure 2). The three residues Arg12 (82), Arg20 (90) and Arg30 (100) are exposed when RALF22 binds to LRX, mediating the interaction with demethylesterified oligogalacturonide (Schoenaers et al., 2024). Overall, the LRX-binding surface of RALF4 and RALF22 is conserved among closely related RALFs, including SlRALF2, SlRALF5 and SlRALF7, but diverges markedly in RALF27, SlRALF9 and SlRALF10 (Figure 2) (Moussu et al., 2023.; Schoenaers et al., 2024). Despite this divergence, these three RALFs retain several positively charged residues at comparable positions (Figure 2). Notably, SlRALF10 contains six acidic amino acids distributed along its sequence, whereas most Arabidopsis RALFs lack acidic residues in their mature sequence, with the exception of RALF4, RALF19 and RALF27, which each contain one or two (Figure 2).

### *Sl*RALF5/7/10 physiological responses in *A. thaliana*

One of the described functions of Arabidopsis RALF peptides is their ability to inhibit cell expansion and growth, as well as to alkalinize the extracellular medium. Most exogenously applied RALF peptides are able to induce root growth inhibition and alkalinization in Arabidopsis suspension cultures (Abarca et al., 2021; Blackburn et al., 2020; Morato do Canto et al., 2014). Here, we screened SlRALF peptides expressed in tomato fruits at higher levels, SlRALF5, SlRALF7 and SlRALF10, for their bioactivity using a root growth inhibition assay in Arabidopsis. Five day-old seedlings were treated with synthetic SlRALF peptides (Supplementary Table S3) for two days before measuring root length. SlRALF5 and SlRALF7 significantly inhibited root growth in three independent biological experiments, whereas SlRALF10 did not (Figure 3a). In a further assay, SlRALF5 caused significant inhibition of seedling growth in Arabidopsis, whereas SlRALF10 showed no effect (Supplemental Figure 2a).

**Figure 3:**
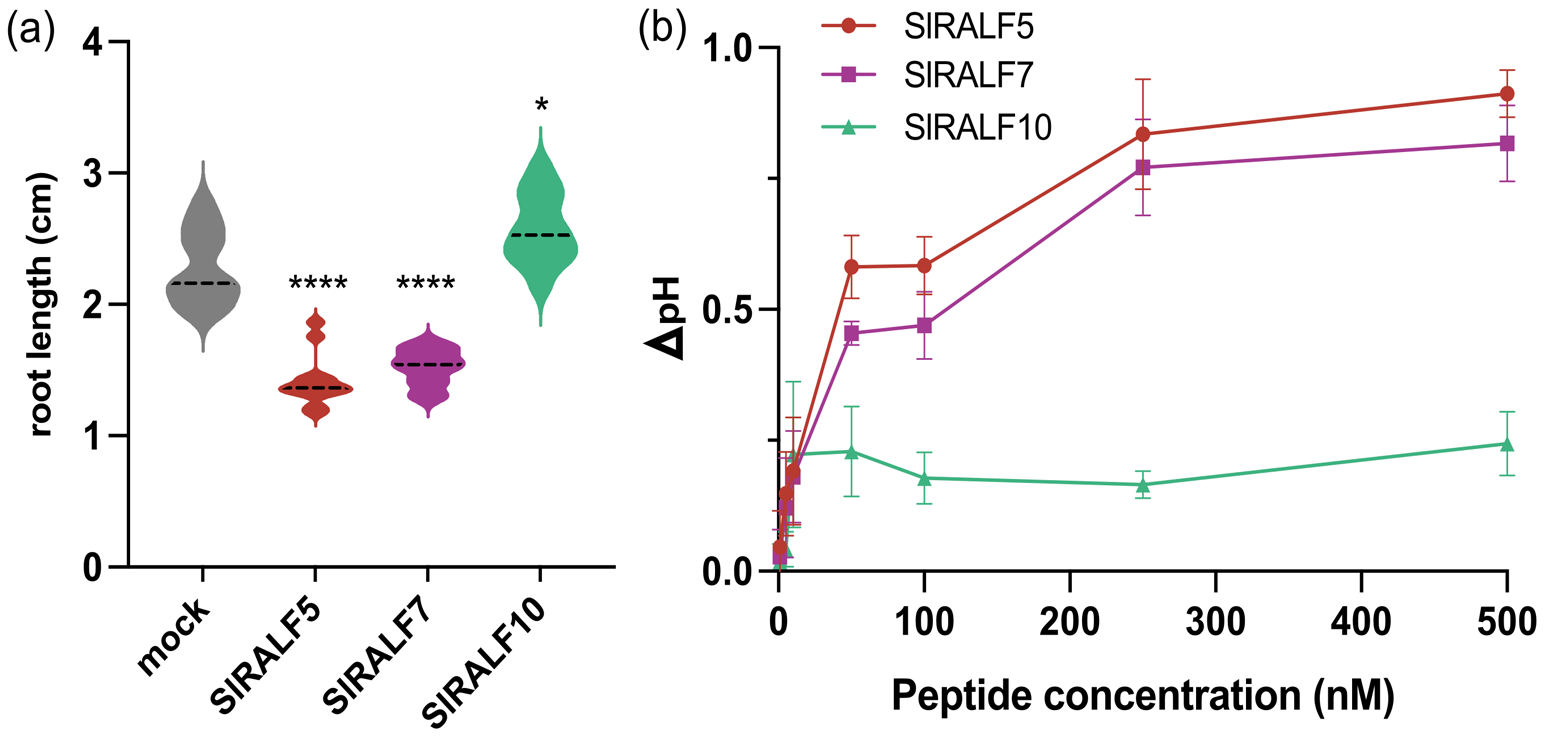
Effect of *Sl*RALF peptides on root growth inhibition and extracellular alkalinization. (a) Primary root length of Arabidopsis 7-d-old seedlings grown in the absence (mock) or presence of 2 μM *Sl*RALF peptides (*n* = 10) Asterisks indicate significance, each treatment was compared with its corresponding mock: *(*P*-value <0,01) and ****(*P*-value <0,0001). (b) Dose-response curves showing ι1pH of tomato cell suspensions after addition of different *Sl*RALF peptides concentration. Data points represent the mean ± standard error of three independent experiments.

We performed alkalinization assays using a customized real-time pH monitoring system to quantitatively measure extracellular alkalinization in tomato cell suspension culture under response to SlRALF peptides (Wang et al., 2024). SlRALF5 and SlRALF7 clearly induced medium alkalinization, while SlRALF10 did not (Figure 3b). Based on these results, we selected SlRALF5 and SlRALF10 for further assays assessing root growth inhibition in tomato. Once again, SlRALF5 caused significant inhibition of root growth, whereas SlRALF10 showed no effect (Supplemental Figure 2b). These results confirm that alkalinization and growth inhibition are coupled characteristics of certain RALF peptides (Abarca et al., 2021). Finally, we investigated whether these tomato fruit RALF peptides trigger reactive oxygen species (ROS) production. Several Arabidopsis RALF peptides have previously been reported to induce significant ROS accumulation (Abarca et al., 2021). None of the SlRALF peptides tested were able to induce a significant ROS response (Supplementary Figure 2c).

### *Sl*RALF5/7/10 interact with *Sl*LRX proteins

In Arabidopsis, several RALFs have been reported to physically interact with LRX and/or CrRLK1L receptor proteins. For example, RALF4 binds to the CrRLK1L proteins BUPS1/2 and ANX1/2, and in a mutually exclusive manner to the LRR domain of LRX8, thereby regulating pollen tube integrity (Ge et al., 2017; Mecchia et al., 2017; Moussu et al., 2023). To examine whether SlRALF5, SlRALF7 and SlRALF10 peptides directly interact with LRX receptor proteins (SlLRX2, SlLRX5), coimmunoprecipitation (Co-IP) assays were performed. The LRR domains of SlLRX proteins were fused to a green fluorescent protein (GFP) tag, and SlRALF peptides were fused to a double hemagglutinin (HA) tag in *Agrobacteirum*-infiltrated *Nicotiana benthamiana* leaves. Immunoprecipitation was conducted using GFP-Trap agarose beads. The three mature SlRALF-HA peptides (SlRALF5, SlRALF7 and SlRALF10) were detected when co-expressed with either SlLRX2-GFP or SlLRX5-GFP (Figure 4). Note that RALF propeptide and mature peptide forms are detected as previously reported (Mecchia et al., 2017).

**Figure 4:**
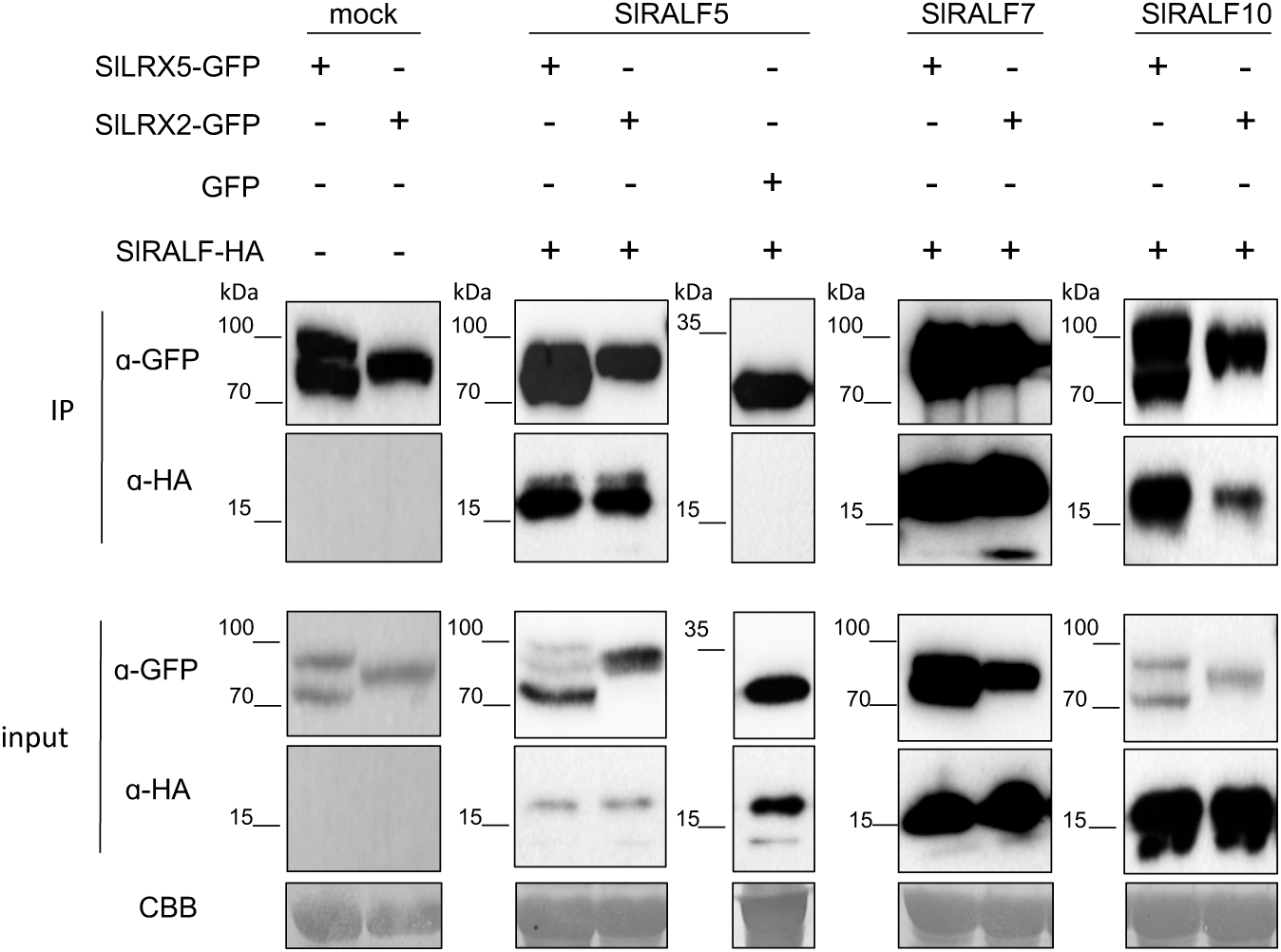
SlRALF5, SlRALF7 and SlRALF10 interact with SlLRX2 and SlLRX5. Western blot analysis of coimmunoprecipitated SlRALF-HA after immunoprecipitation of SlLRX-GFP using antibody against GFP (α-GFP). Predicted sizes: SlRALF5-HA propeptide, 14,4 kDa, and mature 7,5 kDa; SlRALF7-HA propeptide, 16,7 kDa, and mature, 7,9 kDa; SlRALF10-HA propeptide, 12,9 kDa and mature, 7,6 kDa; SlLRX2-GFP, 70,3 kDa; and SlLRX5-GFP, 74,4 kDa. Equal loading was confirmed by Coomassie blue staining (CBB) of input samples. GFP- and HA-tagged proteins were detected with anti-GFP and anti-HA antibody, respectively.

Based on these results, we scrutinized LRR domain sequences of SlLRX2 and SlLRX5 for the presence of sequence motifs previously reported to mediate the interaction of Arabidopsis LRX proteins with RALF peptides (Moussu et al., 2020). To this end, the LRR domain of SlLRX2 was aligned with its closest Arabidopsis homologs, LRX1 and LRX2, which exhibited 63,1% and 64,8% sequence identity, respectively. Similarly, the LRR domain of SlLRX5 was aligned with LRX3, LRX4 and LRX5, showing 72,6%, 72,4% and 69,2% sequence identity, respectively (Supplementary Figure 3). All five pairs of Cys residues previously described to form intra-molecular disulfide bonds, as well as the Cys residue responsible for the inter-molecular disulfide bridge covalently linking the two LRX protomers in Arabidopsis LRXs, are fully conserved in SlLRX2 and SlLRX5 (Supplementary Figure 3) (Moussu et al., 2020). We subsequently analyzed in detail the amino acids residues constituting the LRX-RALF4 binding pocket (Moussu et al., 2020). Within the described secondary structure, particularly in the β3 and β5 loops located between the first Cys dimer and the inter-protomer Cys dimer, SlLRX5 displays an enrichment in polar residues relative to most Arabidopsis LRXs, with the exception of LRX7, which also exhibits a similar enrichment (Supplementary Figure 3) (Moussu et al., 2020). Overall, the high conservation of the RALF-binding pocket suggests that SlLRX2 and SlLRX5 may interact with RALF peptides in a conformation analogous to that characterized for Arabidopsis LRXs (Moussu et al., 2020).

### Prediction of the SlRALF5/10 mature peptide interaction with SlLRX5

Next, we modelled the SlLRX5-SlRALF5 and SlLRX5-SlRALF10 complexes using AlphaFold2, including the LRR domain and the mature SlRALF peptides (Figure 5). SlLRX5 is structurally related to the LRX proteins LRX2 and LRX8 described in Arabidopsis (Lee & Santiago, 2023; Moussu et al., 2020). Consistent with experimentally determined LRX-RALF structures, SlRALF peptides adopt a folded, disulfide bond-stabilized conformation that fits into the conserved binding pocket within the LRR domain through extensive contacts. Our model predicts that SlRALF5 binds SlLRX5 in a configuration compatible with that described for Arabidopsis LRX-RALF. In this orientation, residues corresponding to R82, R90 and R100 of Arabidopsis RALF22 align with K78, R85 and K94 in SlRALF5 and remain solvent-exposed, consistent with the polycationic pectin-interaction surface in Arabidopsis (Figure 5a) (Moussu et al., 2020; Schoenaers et al., 2024). In contrast, the predicted SlLRX5-SlRALF10 complex displays a distinct peptide conformation within the binding pocket. Although residues corresponding to the conserved polycationic surface (K62, K69 and K78 in SlRALF10) are present, their proximity to a negatively charged residue is predicted to alter their orientation and potentially modify the electrostatic surface of the peptide (Figure 5b).

**Figure 5:**
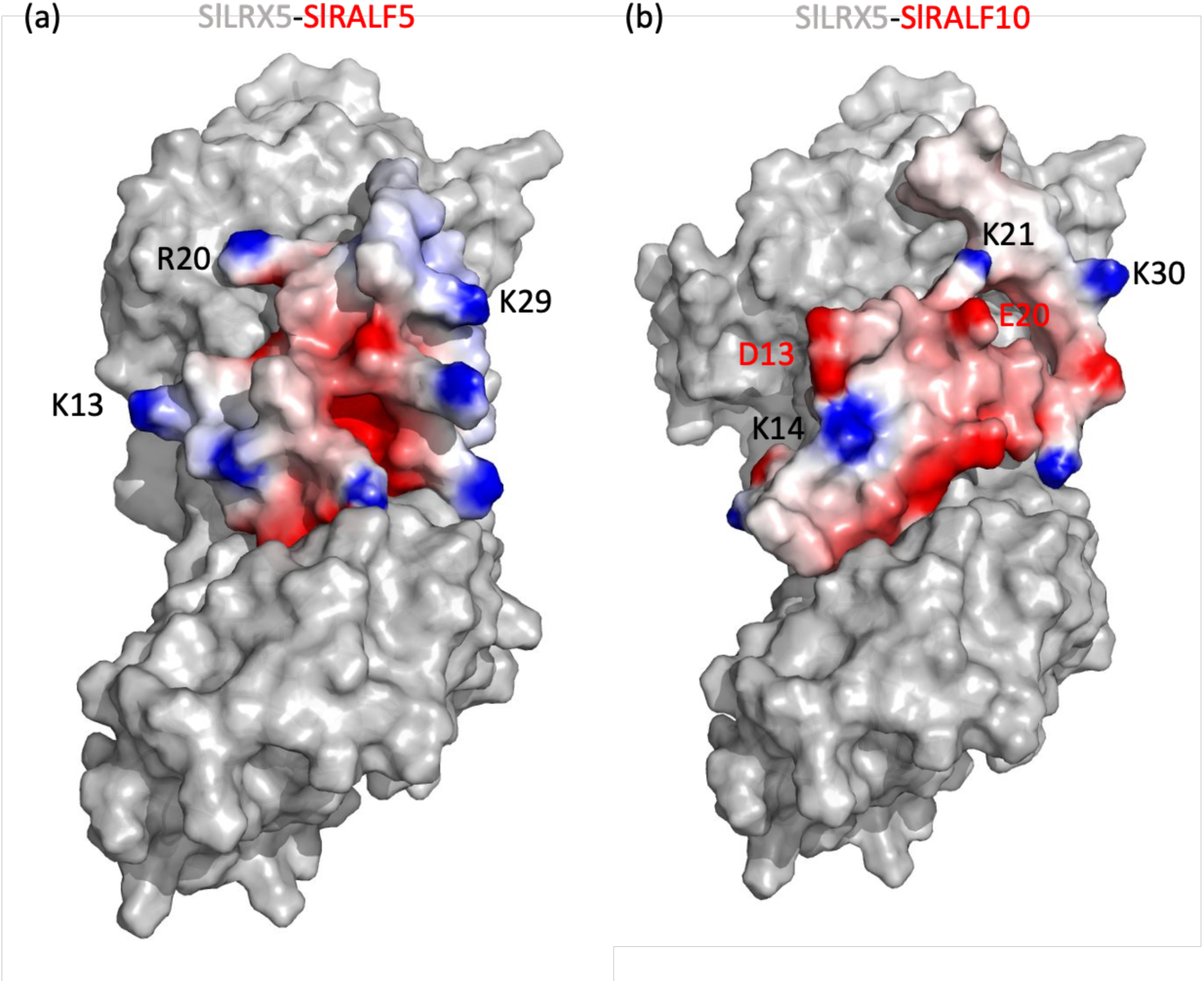
Exposed surface of SlRALF5 and SlRALF10 when bound to SllRX5. Electrostatic surface representation of SlRALF5 (a) and SlRALF10 (b) bound to SlLRX5. Solven-accessible surface electrostatic potential has been calculated using the adaptative Poisson-Boltzmann solver (APBS) plugin (PyMOL) at pH 5. The potential is given with the negative (red) and positive (blue) contour levels in the range from -5.0 to +5.0 kBT. Details of the free residues are shown.

## Discussion

RALF peptides have been extensively studied in root development, plant reproduction, abiotic stress responses, and immunity (Liu et al., 2024; Lu et al., 2025). In contrast, their roles during fruit development and ripening remain poorly understood. Here, we provide an initial framework linking SlRALF peptides to tomato fruit biology. Tissue-specific expression analyses revealed three major expression patterns among *SlRALF* genes (Figure 1a). SlRALF1, SlRALF3, SlRALF6, and SlRALF8 are predominantly expressed in floral tissues, whereas SlRALF2 is mainly expressed in roots. By contrast, SlRALF5, SlRALF7, SlRALF9, and SlRALF10 show broader expression profiles. In fruit tissues, these four genes are expressed at the mature green stage, with SlRALF10 displaying the highest transcript abundance. However, only SlRALF5 and SlRALF7 remain expressed throughout ripening, suggesting specialized functions during late fruit development.

Phylogenetic analyses further support functional diversification within this family. SlRALF2, SlRALF5, and SlRALF7 cluster with Arabidopsis RALF1/RALF22/RALF23/RALF33, whereas SlRALF9 and SlRALF10 group with Arabidopsis RALF27 (Montano et al., 2026). Because RALF27 remains functionally uncharacterized, the analysis of SlRALF10 may also provide insights into the role of this poorly understood clade. Structural analyses revealed substantial differences between SlRALF10 and SlRALF2/5/7. Most SlRALF peptides are enriched in basic amino acids, but SlRALF2 and SlRALF10 are notable exceptions, displaying acidic pI values (Wang et al., 2025). SlRALF10 also shows a markedly different charge distribution compared with canonical RALFs such as Arabidopsis RALF4 (Figure 2) (Moussu et al., 2023). In addition, motifs associated with LLG interaction are highly conserved in SlRALF2/5/7 but only partially retained in SlRALF10. The GASYY and PYXRGCS motifs are similarly conserved in SlRALF2/5/7 but diverge in SlRALF9/10. Together, these observations suggest that SlRALF10 interacts with extracellular partners through mechanisms distinct from those of SlRALF2/5/7.

Differences in putative LRX interactions further support this idea. Two conserved Tyr residues stabilize RALF binding to the LRX pocket (Moussu et al., 2020), yet SlRALF10 retains only one of them (Figure 2). This feature may weaken LRX association and favor a more dynamic equilibrium between LRX-bound and LLG–FER-associated states. Moreover, residues involved in pectin interaction are strongly conserved in SlRALF2/5/7 but not in SlRALF10. In Arabidopsis, RALF4 and RALF22 interact electrostatically with demethylesterified pectin through positively charged surfaces (Moussu et al., 2023; Schoenaers et al., 2024). By contrast, SlRALF10 contains negatively charged residues within the predicted interaction interface, which may alter its affinity for pectin or favor binding to specific methylesterification status.

These structural differences support a model in which distinct SlRALF peptides could respond differently to ripening-associated changes in the apoplast. During tomato ripening, pectin demethylesterification and degradation progressively increase the abundance of negatively charged oligogalacturonides (OGs) (Wang & Seymour, 2022). Because SlRALF5/7 retain highly conserved positively charged surfaces, these peptides may become increasingly sequestered by OGs through electrostatic interactions. Such sequestration could reduce their availability for CrRLK1L receptor binding and attenuate downstream signaling. In contrast, SlRALF10 may operate through a more dynamic interaction regime that depends on pH, cell wall status, and the balance between LRX-and LLG/CrRLK1L-associated complexes. This functional diversification may provide a mechanism linking cell wall integrity with receptor-mediated signaling during fruit ripening. These contrasting structural properties of SlRALFs are consistent with their distinct biological activities. SlRALF5/7 inhibited root growth and induced alkalinization of the growth medium, whereas SlRALF10 did not (Figure 3 and Supplementary Figure 2). Similar phenotypes have been reported for Arabidopsis RALF27, which also lacks growth inhibitory and alkalinization activity (Abarca et al., 2021). These observations support the idea that SlRALF10 belongs to a functionally distinct subgroup. Although several RALF peptides induce ROS accumulation (Abarca et al., 2021), neither SlRALF5 nor SlRALF10 triggered significant ROS production under our experimental conditions (Supplementary Figure 2).

SlCrRLK1L20, the closest tomato homolog of Arabidopsis FER, is highly expressed during fruit ripening and positively regulates both ripening progression and resistance to *Botrytis cinerea* (Ji et al., 2020, 2023; Ma et al., 2023). Because SlRALF5 and SlRALF7 interact with SlCrRLK1L20ECD in yeast two-hybrid assays (Wang et al., 2025), these peptides may function upstream of FER-like signaling pathways during ripening. This raises the possibility that RALF-mediated sensing of cell wall status contributes to the coordination of ethylene signaling, defense responses, and fruit softening. In parallel, our results identify LRX proteins as additional components of this regulatory system. SlLRX5 and, to a lesser extent, SlLRX2 are expressed throughout fruit development and temporally overlap with SlRALF5/7/10 expression (Figure 1b). Co-IP assays confirmed interactions between these peptides and both SlLRX proteins (Figure 4), supporting the existence of LRX–RALF complexes in tomato fruit. Among them, SlLRX5 likely plays the predominant role. Its interaction with SlRALF10 may be especially relevant during early fruit development, whereas SlRALF5/7 interactions may predominate during ripening.

Structural modeling further supports differential interaction modes among SlRALFs. The predicted SlLRX5–SlRALF5 complex closely resembles experimentally characterized Arabidopsis RALF–LRX structures and retains a positively charged binding surface compatible with pectin interaction (Figure 5A). By contrast, the predicted SlLRX5–SlRALF10 complex displays a less positively charged interface due to the presence of exposed acidic residues (Figure 5B). Although these predictions require experimental validation, they support the idea that distinct SlRALF peptides could engage cell wall components through different electrostatic interaction patterns. These observations fit well with the extensive cell wall remodeling that accompanies fruit ripening. Pectin depolymerization and demethylesterification progressively weaken the wall and increase apoplastic porosity (Posé et al., 2019; Wang & Seymour, 2022). PME activity initially generates demethylesterified homogalacturonan domains that can transiently reinforce the wall through Ca²⁺-mediated crosslinking, but subsequent degradation by pectate lyase and polygalacturonase ultimately drives irreversible softening (Peng et al., 2022; Sénéchal et al., 2014; Uluisik & Seymour, 2020). Within this dynamic environment, RALF–LRX–FER modules may act as sensors that translate changes in pectin status and apoplastic pH into signaling outputs that coordinate wall remodeling, ripening progression, and defense responses. Collectively, our results provide a theoretical model in which distinct SlRALF peptides differentially may signal ripening-associated cell wall remodeling through interactions with LRX and CrRLK1L proteins. Further functional studies will clarify how this signaling network is linked to pectin status, apoplastic pH, and receptor-mediated pathways to coordinate fruit softening during tomato ripening.

## Materials and Methods

### Gene expression pattern and phylogenetic analyses

The *SlRALF* and *SlLRX* gene expression pattern analyses used RNA-seq data from the SGN tomato functional genomic database (SGN-TFGD, http://ted.bti.cornell.edu/cgi-bin/TFGD/digital/home.cgi) (Sato et al., 2012). The data are listed in Tables S1 and S2. The heatmap was illustrated with Morpheus (https://software.broadinstitute.org/morpheus/). The RALF and LRX proteins sequences from tomato and Arabidopsis were downloaded from the National Center for Biotechnology Information (NCBI, https://www.ncbi.nlm.nih.gov/) and TAIR (https://www.arabidopsis.org/), respectively. The obtained full protein sequences were aligned by ClustalW to construct a phylogenetic tree with the default parameters. Multiple sequence alignment of the mature peptide sequence were created using ClustalW.

### Plant material and growth conditions

We used *A. thaliana* (Col-0 ecotype), *S. lycopersicum* (Money Maker) and *N. benthamiana* as experimental materials. Arabidopsis and tomato seeds were surface sterilized with chlorine vapors (100 mL commercial bleach + 3 mL 37% HCl) in a sealed container for 3-4h, then air-cleared for 2-4h in a laminar flow cabinet. Seeds were plated under sterile conditions on ½ Murashige-Skoog medium with 1% (w/v) sucrose and 1% (w/v) agar. Plated seeds underwent 3-d vernalization at 4°C in darkness, then grew vertically in a chamber (16h light/8h dark, 130 +/- 30μmol photons m^-2^ s^-1^, 22 +/- 1°C).

### Peptides

SlRALF peptides were synthesized (Supplemental Table S3) by GenScript Biotech (www.genscript.com) with a purity of >90%. The peptides were dissolved in sterile pure water for usage and stored at -20°C at a concentration of 1 mM.

### Root growth inhibition assay

Arabidopsis and tomato seeds were grown for 5 d before transferring ten seedlings to each well of a 12-well plate containing 4 mL per well of ½ MS medium containing 2 μM RALF. Seedlings were transferred 2 d later to solid ½ MS plates. Control seedlings were performed under identical conditions in a peptide-free medium. Primary root length was measured by scanning the plates and quantified using the software Fiji (https://imagej.net/Fiji). Experiments were repeated at least three times using independent biological replicates.

### Seedling growth inhibition assay

Arabidopsis seeds were grown for 5 d before transferring ten seedlings to each well of a 12-well plate containing 4 mL per well of ½ MS medium containing 2 μM RALF. Seedlings weight was measured 7 d later. Control seedlings were performed under identical conditions in a peptide-free medium. Experiments were repeated at least three times using independent biological replicates.

### Alkalinization assay

Extracellular alkalinization assays were performed in tomato (*Solanum peruvianum*) wild type suspension cultures as described (Wang et al., 2024; Li et al., 2025). Briefly, RALF peptides were diluted and added into 10 mL of seven-day old cell suspension from 200-fold concentrated stock solutions prepared in ddH2O. Following elicitor treatment, pH of treated cells was recorded at 10s intervals for 60 mins. Dosage curve was then plotted using peak values at each concentration. At least three biological replicates were performed for each treatment.

### ROS production assay

Arabidopsis seedlings were sterilized and vernalized as described above. Plates were transferred to short day photoperiod (8h light/16h dark) for 4-5 days. After this period, plants were transferred to soil composed of organic substrate and vermiculite (4:1 v/v) during four weeks under short day. ROS measurement was performed as previously described (Sang and Macho, 2017). Plant discs were incubated overnight with water. The next day, water was replaced by 75 μL 2 mM MES-KOH pH 5,8 to mimic the apoplastic pH. Leaf discs were incubated for 4h before adding 75 μL 40 μg/mL horseradish peroxidase (HRP), 1 μM L-O12 (Wako Chemicals) and 4 μM RALF peptide. In total, 24 leaf discs were used for each condition. Luminescence was measured using a GloMax 96 Microplate Luminometer (Promega).

### Plasmid constructs

*SlLRX2* and *SlLRX5* were amplified from *S. lycopersicum* genomic DNA with high-fidelity DNA polymerase (Q5 High-Fidelity DNA Polymerase, New England Biolabs, #M0491). *SlLRX* genes were cloned until the Cys-rich region end predicted in (Moussu S et al., 2020). All the primer sequences are listed in Supplementary Table S4. Using Multisite Gateway cloning, the PCR product were inserted into pDONR L1-L2 via BP reactions (Invitrogen). All constructs were verified by PCR, restriction analysis and sequencing. The sequences encoding SlRALF peptides, together with sequenced coding for a 2xHA-epitope tag flanked by att sites (Gateway, Life Technologies), were synthesized using Genscript technology (Genscript, NJ, USA): SlRALF5 (5’ CAAATAATGATTTTATTTTGACTGATAGTGACCTGTTCGTTGCAACACATTGATGAGCAATGCT TTTTTATAATGCCAACTTTGTACAAAAAAGCAGGCTCCATGGGAGTTTCTTCGTATTTGATTGTTTGTGTTCTTGTTGGAGCTTTTTTCATTTCCATGGCTGCCGCCGGCGACAGTGGTAGCTACGATTGGATGGTGCCGGCGAGATCCGGTGAATGCAAGGGGAGTATTGCAGAGTGCATGGCTGAAGAAGATGAGTTTGCCCTTGACAGTGAGTCAAACAGGCGTATTTTAGCAACCAAAAAGTACATCAGCTATGGTGCACTCCAGAAGAACAGTGTGCCGTGTTCTCGCCGTGGAGCTTCCTACTACAACTGCAAACCTGGAGCTCAAGCAAATCCCTACACTCGTGGATGCAGTGCTATTACTCGTTGCAGGAGCTACCCATACGATGTTCCAGATTACGCTTACCCATACGATGTTCCAGATTACGCTTAGCACCCAGCTTTCTTGTACAAAGTTGGCATTATAAGAAAGCATTGCTTATCAATTTGTTGCAACGAACAGGTCACTATCAGTCAAAATAAAATCATTATTTG3’), SlRALF7 (5’CAAATAATGATTTTATTTTGACTGATAGTGACCTGTTCGTTGCAACACATTGATGAGCAATGCTTTTTTATAATGCCAACTTTGTACAAAAAAGCAGGCTCCATGGCGAAAGTGAAATCTCTCTCTATTTTCTTCATCTCATCCATTTTCTCCGTCGTTATTGTAGCGCTTTTATCTCCGGCCGCCGCTGGTTCCGCGGCGGCAGTAGCTGCTACTGGGTCCCACCAGCTGAGTTACTTTCCGATGACATTATCTTCTTCTTCTTCACCGATCTGCGACGGTTCGATCGGGGACTGTTTGGCTGAAGAAGACGAGAATGAGTTCGGGATGGAATCGGAGAGCAGCCGGCGCATGTTAGCATACCGCCGGAGATACATTAGCTACGGCGCGCTTAGTAGTAACAGAGTGCCGTGTTCAAGGAGAGGAGCTTCGTACTACAATTGCCGTCCCGGAGCTCAGGCGAATCCTTATCAACGTGGATGCAGTGCCATCACGCGCTGCCGTCATTACCCATACGATGTTCCAGATTACGCTTACCCATACGATGTTCCAGATTACGCTTAGCACCCAGCTTTCTTGTACAAAGTTGGCATTATAAGAAAGCATTGCTTATCAATTTGTTGCAACGAACAGGTCACTATCAGTCAAAATAAAATCATTATTTG3’) and SlRALF10 (5’ CAAATAATGATTTTATTTTGACTGATAGTGACCTGTTCGTTGCAACACATTGATGAGCAATGCTTTTTTATAATGCCAACTTTGTACAAAAAAGCAGGCTCCATGAGGAGCTGCTGCAGTTTGTTTTGGCTATCCATGGTGATTTTGCTCTCTGCTTCTTCGGTAACAGACGGGACGATGATCGTCGAAAGGCGGTGGAACGGCACCGGAGAAATGGATGGTATTGATTGGGAGGTTAGCCTTGCAGGTGATGTAAATTACTTGTCTTACAAAGTTACTAATGATAAGCCACCGATCTGCAACGAGAAAGTGTACGGGAATTGCATAGGGGACAAGAAGCCTGGCCAACGCCCATGTACTGAATACAATAACTGCAAGCGTTCTCCAGATTACCCATACGATGTTCCAGATTACGCTTACCCATACGATGTTCCAGATTACGCTTAGCACCCAGCTTTCTTGTACAAAGTTGGCATTATAAGAAAGCATTGCTTATCAATTTGTTGCAACGAACAGGTCACTATCAGTCAAAATAAAATCATTATTTG3’).

For expression plasmid generation, the entry vectors (pENTRY L1-Gene-L2) were recombined with destination vectors using a LR Clonase^TM^ reaction (Invitrogen). Specifically, pGWB5 was used for receptor constructs and pGWB2 for peptide constructs. The resulting plasmids are 35S:SlLRX2-GFP, 35S:SlLRX5-GFP; and 35S:SlRALF5-HA, 35S:SlRALF7-HA and 35S:SlRALF10-HA.

### Transient expression in *N. benthamiana*

The binary plasmids were transformed into A. tumefaciens strain GV3101::pMP90 by electroporation. *A. tumefaciens* strains were grown overnight at 28°C in LB medium with rifampicin (50 μg/mL), gentamicin (25 μg/mL), and the construct specific antibiotic kanamycin (50 μg/mL). Cultures were centrifuged (3,000 x g, 15 min, RT), and pellets were resuspended in agroinfiltration solution (10 mM MES pH5.6, 10 mM MgCl2, 1 mM acetosyringone), then incubated in the dark for 2-4h at RT. For double-gene expression, Agrobacterium suspensions were mixed to reach OD_600_ of 0.33 per constructs and 0.25 for p19, to prevent gene silencing. Two leaves from 3-week-old *N. benthamiana* plants (3^rd^ and 4^th^ from the apex) were infiltrated on the abaxial side using a needleless 1 mL syringe. Plants were maintained under growth conditions for 2 d before sample collection.

### Co-immunoprecipitation assay and immunoblot analysis

Co-IP experiments were conducted based on the methodology described in (Amorim-Silva et al., 2019). *N. benthamiana* leaves transiently transformed were ground in liquid nitrogen to a fine powder. Proteins were extracted from 0.5 g of tissue by incubating the ground material for 30 min at 4°C under rotation with 1 mL of nondenaturing extraction buffer (150 mM Tris-HCl pH 7.5, 150 mM NaCl, 1% (v/v) Nonidet P-40, 10 mM EDTA, 1 mM Na_2_MoO_4_, 1 mM NaF, 10 mM DTT, 0.5 mM Pefabloc^®^, 1% (v/v) protease inhibitor cocktail (Sigma, #P9599)). The resulting extracts were centrifuged (16,000 x g, 20 min, 4°C) and filtered through Poly-Prep chromatography columns (Bio-Rad, #731-1550). An aliquot of 100 μL was set aside as input for Western blot analysis, while the remaining extract was incubated for 2h with 30 μL GFP-Trap^®^ Magnetic Agarose beads (Chromotek). To reduce unspecific interactions, Nonidet P-40 concentration was adjusted to 0.2% during incubation. Beads were washed four times with detergent-free extraction buffer, then resuspended in 75 μL of 2 x Laemmli Buffer (125 mM Tris-HCl pH6.8, 4% (w/v) SDS, 20% (v/v) glycerol, 2% (v/v) β-mercaptoethanol, 0.01% (w/v) bromophenol blue). Samples were heated at 95°C for 5 min, and subsequently centrifuged (20,000 x g, 1 min RT) to dissociated immunocomplexes. Proteins extracts were separated by molecular weight using 10% SDS-PAGE. Electrophoresis was carried out at a constant voltage of 80 V during stacking, followed by 120 V for protein separation. Subsequently, proteins were transferred onto a methanol-equilibrated PVDF membrane (Immobilon-P, Millipore, 0.45 μm, #IPVH00010) using a semi-dry transfer method following the standard protocol of the Trans-Blot Turbo Transfer System (Bio-Rad), at 25 V for 30 min. Membranes were then blocked with 5% (w/v) skimmed milk in Tris-buffered saline containing 0.05% Tween-20 (TTBS) (1h, RT). After blocking, membranes were washed and incubated overnight at 4°C with gentle agitation in the presence of specific primary antibodies: mouse anti-GFP 1:4000 (Santa Cruz Biotechnology, #sc-9996) or mouse anti-HA 1:3000 (Sigma-Aldrich, #H3663). After washing, membranes were incubated (2h, RT) with HRP-conjugated secondary antibodies anti-mouse 1:80000 (Sigma-Aldrich, #A9044). Signal detection was carried out by chemiluminescence using a ChemiDoc XRS+ system (BioRad) with either Clarity ECL Western Blotting Substrate (BioRad, #170-5060) or SuperSignal West Atto Ultimate Sensitivity Substrate (Thermo Scientific, #A38555). Exposure times varied from a few seconds up to 10 min, ensuring only unsaturated images were analyzed. After immunodetection, membranes were stained with Coomassie R-250 to verify equal protein loading. Input and IP samples were run in duplicate SDS-PAGE gels for GFP and HA detection. Each Co-IP assay was repeated three times with consistent results.

### AlphaFold structural prediction and analysis

Structural models of the SlLRX5^LRR^–SlRALF5 and SlLRX5^LRR^–SlRALF10 complexes were generated using the AlphaFold 3 server (https://alphafoldserver.com) (Abramson et al., 2024). The full-length SlLRX5^LRR^ sequence and the mature peptide sequences of SlRALF5 (ATKKYISYGALQKNSVPCSRRGASYYNCKPGAQANPYTRGCSAITRCRS) and SlRALF10 (GDVNYLSYKVTNDKPPICNEKVYGNCIGDKKPGQRPCTEYNNCKRSPD) were submitted as separate protein chain inputs. Five structural models were generated per complex; the top-ranked model (model_0) was selected based on the highest per-residue confidence score (pLDDT) and interface predicted TM-score (ipTM). Models with ipTM > 0.6 were considered reliable predictions of the protein–protein interaction interface. For comparison, the ColabFold implementation of AlphaFold2-Multimer (alphafold2_multimer_v3) was also used as an alternative approach. Structural superposition of the tomato complexes with the reference LRX2–RALF4 structure (PDB: 6QXP) (Moussu et al., 2020) was performed by aligning the LRR domains (chain A) of each model using PyMOL (The PyMOL Molecular Graphics System, version 2.5, Schrödinger). Electrostatic surface potential was calculated using the APBS (Adaptive Poisson-Boltzmann Solver) plugin for PyMOL at pH 5.0 to simulate apoplastic conditions, with a solvent ion concentration of 0.15 M KCl and a grid spacing of 0.5 Å. The electrostatic potential was mapped onto the RALF peptide surface using a colour scale from −5 to +5 k_B_T/e (red, negative; blue, positive), as previously described (Lee & Santiago, 2023; Moussu et al., 2020). All structural figures were rendered with PyMOL.

### Statistic

Graphs and statistical analyses were performed using Prism 8.02 (GraphPad Software, www.graphpad.com), by applying one-way ANOVA followed by Dunnett’s multiple test (****P<0,0001; *P<0,01). Asterisks in figures indicate statistical differences between peptide treatments and mock. Experiments were repeated at least three times with similar results.

## Acknowledgements

This work was funded by grant PID2020-113378RA-I100 (Ministerio de Ciencia e Innovación, Spain) to JAM, PMR and VGD; RYC2018-024032 (Ministerio de Ciencia e Innovación, Spain) to AML and VGD; and UMA20JA-FEDERJA-055 (Junta de Andalucía and Universidad de Málaga, Spain) to MC and VGD.

## Author contributions

Conceptualization: VGD; Investigation: JAM, MC, XW, PMR, AML, AS, IM, VGD; Writing-original draft: JAM, VGD; Writing- review & editing: all authors.

## Supporting information

**Supplementary Figure 1:**
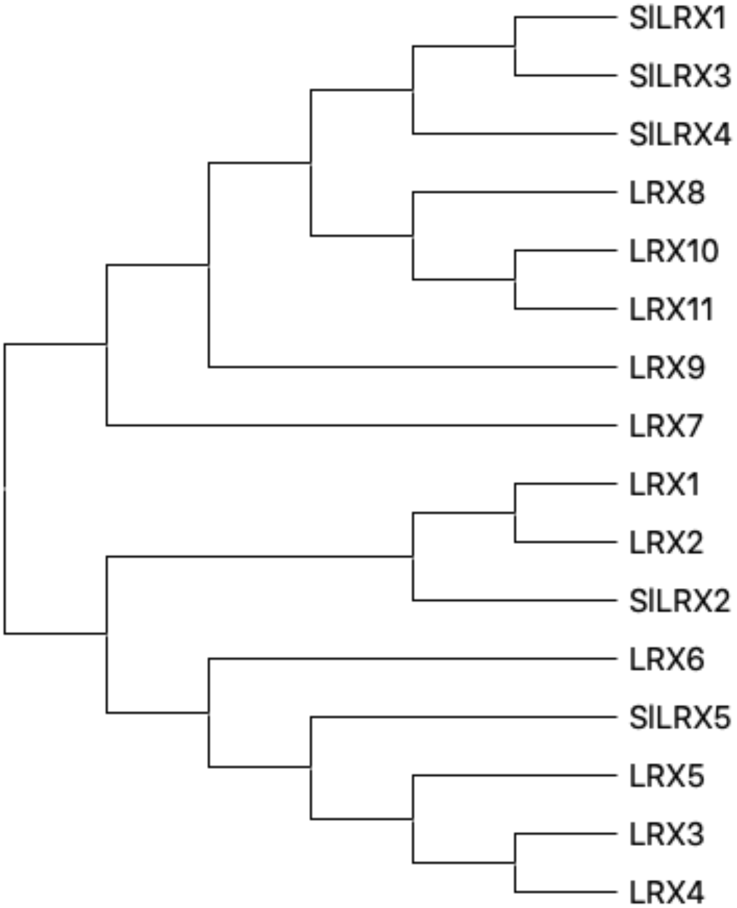
Phylogenetic tree of LRX proteins based on an alignment of tomato and Arabidopsis sequences. The unrooted phylogenetic tree was constructed with ClustalOmega using the neighbor-joining method with 1000 bootstrap replicates and implemented in MEGA 5.0.

**Supplementary Figure 2:**
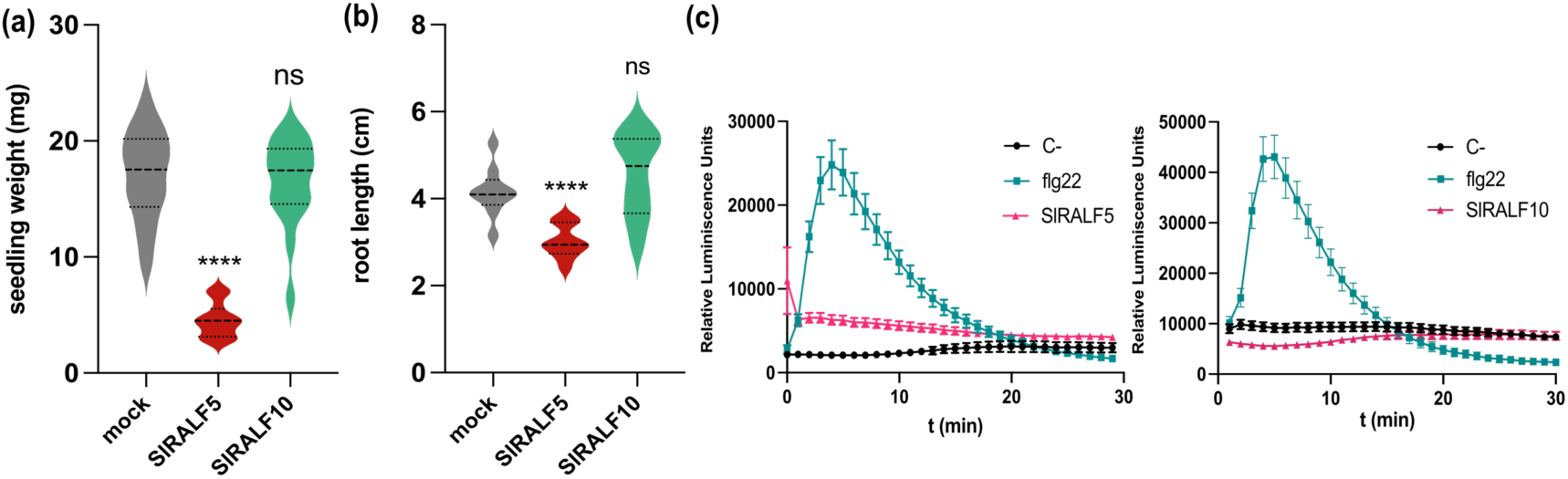
Effect of *Sl*RALF peptides on seedling weight, root growth inhibition and ROS production. (a) Fresh weight of 12-d-old Arabidopsis seedlings grown in the absence (mock) or presence of 2 μM SlRALF peptide (*n* = 10). (b) Primary root length of tomato 7-d-old seedlings grown in the absence (mock) or presence of 2 μM *Sl*RALF peptides (*n* = 10) Asterisks indicate significance, each treatment was compared with its corresponding mock: ns (*P*-value >0,05) and ****(*P*-value <0,0001). (c) ROS production in Arabidopsis leaf discs treated with 4 μM SlRALF peptide. Shown are the kinetics of one representative replicate.

**Supplementary Figure 3:**
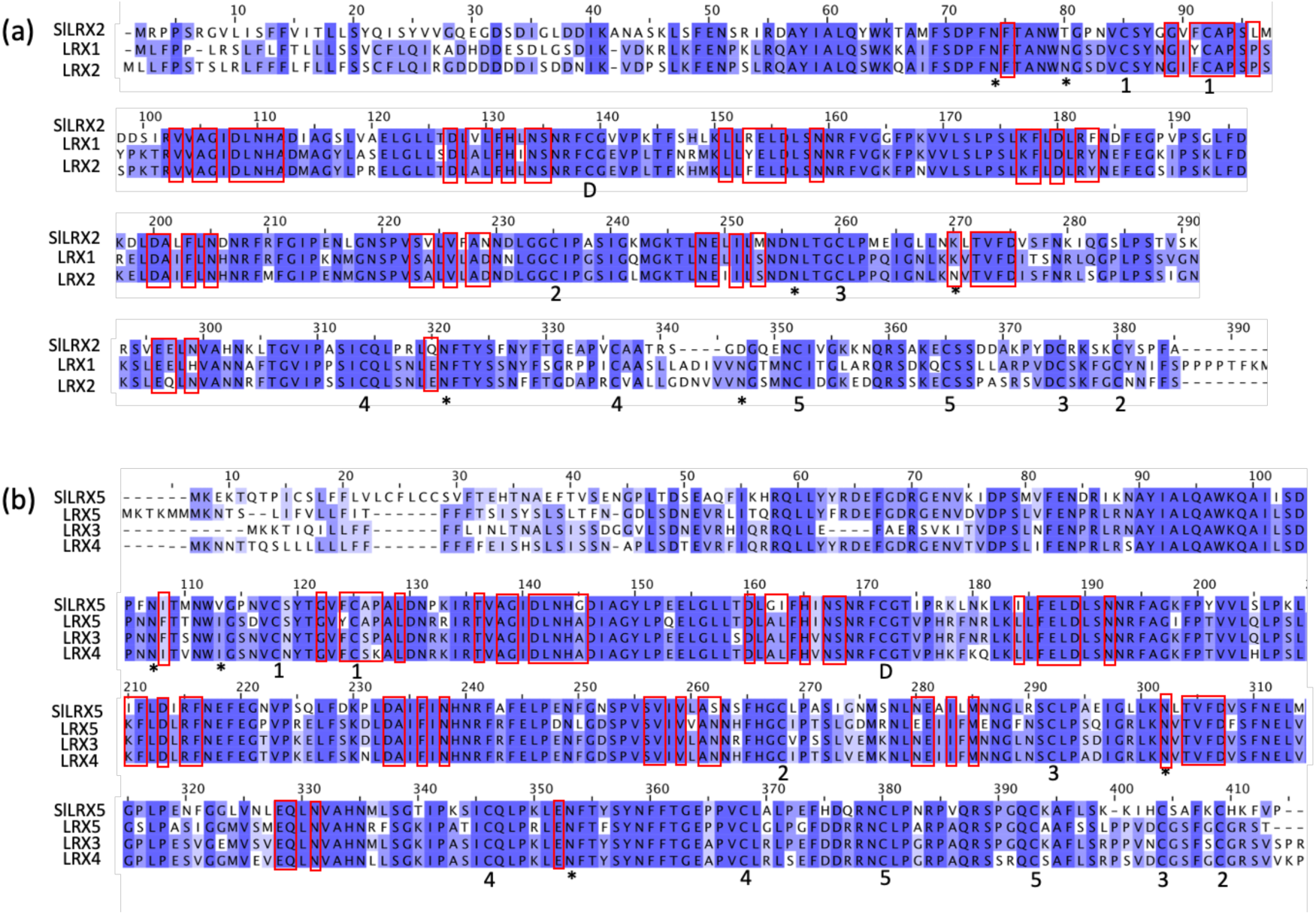
Sequence alignment of the LRR domain of SlLRX2 (a) and SlLRX5 (b) with their related Arabidopsis LRX members. Conserved residues are highlighted in different blue tones. Motifs reported to mediate interactions with RALF forming the binding pocket are enclosed in red boxes. Conserved Cys residues engaged in disulfide bonds with matching pairs are indicated by numbers. N-glycosylation sites are indicated by black stars.

**Table S1:** Expression data of *SlRALF* in different tissues.

| Gene | Expression data from SGN-TFGB (RPKM) |  |  |  |  |  |  |  |  |  |
| --- | --- | --- | --- | --- | --- | --- | --- | --- | --- | --- |
|  | Heinz_root | Heinz_leaf | Heinz_bud | Heinz_flower | Heinz_1cm_fr | Heinz_2cm_fr | Heinz_3cm_fr | Heinz_MG | Heinz_Br | Heinz_Br+10 |
| *SIRALF1* | 0.23 | 0.22 | 119.31 | 232.54 | 0.54 | 0 | 0.22 | 0 | 0 | 0 |
| *SIRALF2* | 417.99 | 1.37 | 3.38 | 5.64 | 1.34 | 1.14 | 0.83 | 2.3 | 1.32 | 1.74 |
| *SIRALF3* | 0.13 | 0 | 103.25 | 52.12 | 0.31 | 0 | 0 | 0.34 | 0 | 0 |
| *SIRALF4* | 6.25 | 9.26 | 6.4 | 10.11 | 20.89 | 24.37 | 13.59 | 12.15 | 5.04 | 0.59 |
| *SIRALF5* | 146.86 | 104.2 | 82.15 | 89.11 | 167.56 | 65.96 | 31.66 | 165.29 | 125.5 | 175.86 |
| *SIRALF6* | 0.66 | 0 | 289.73 | 559.44 | 0.77 | 0.19 | 1.02 | 0.79 | 0.13 | 0 |
| *SIRALF7* | 34.36 | 61.62 | 93.7 | 164.88 | 101.63 | 95.79 | 45 | 64.33 | 67.8 | 65.84 |
| *SIRALF8* | 0 | 18.08 | 103.27 | 6.13 | 88.32 | 32.46 | 32.87 | 2.34 | 0.4 | 0 |
| *SIRALF9* | 4.28 | 260.64 | 203.76 | 106.08 | 164.89 | 89.1 | 70.53 | 56.62 | 4.96 | 0.99 |
| *SIRALF10* | 331.78 | 178.03 | 191.55 | 145.57 | 396.55 | 271.85 | 319.43 | 326.12 | 26.37 | 1.33 |
| *SIRALF11* | 0 | 0 | 0 | 0 | 0 | 0 | 0 | 0 | 0 | 0 |

**Table S2:** Expression data of *SlLRX* in different tissues.

| Gene | Expression data from SGN-TFGB (RPKM) |  |  |  |  |  |  |  |  |  |
| --- | --- | --- | --- | --- | --- | --- | --- | --- | --- | --- |
|  | Heinz_root | Heinz_leaf | Heinz_bud | Heinz_flower | Heinz_1cm_fr | Heinz_2cm_fr | Heinz_3cm_fr | Heinz_MG | Heinz_Br | Heinz_Br+10 |
| *SILRX1* | 0.14 | 0 | 20.1 | 75.17 | 0.09 | 0.03 | 0.35 | 0.23 | 0.11 | 0 |
| *SILRX2* | 95.48 | 2.18 | 7.13 | 3.73 | 49.82 | 81.44 | 49.82 | 80.32 | 43.51 | 57.3 |
| *SILRX3* | 1.1 | 0.21 | 77.98 | 400.34 | 0.28 | 0.06 | 1.62 | 0.82 | 0.35 | 0 |
| *SILRX4* | 0.7 | 0.37 | 4.57 | 18.05 | 0.3 | 0.37 | 0.18 | 0.37 | 0.55 | 0.96 |
| *SILRX5* | 93.17 | 23.79 | 20.8 | 25.12 | 42.37 | 89.77 | 57.94 | 322.71 | 471.42 | 825.65 |

**Table S3:**
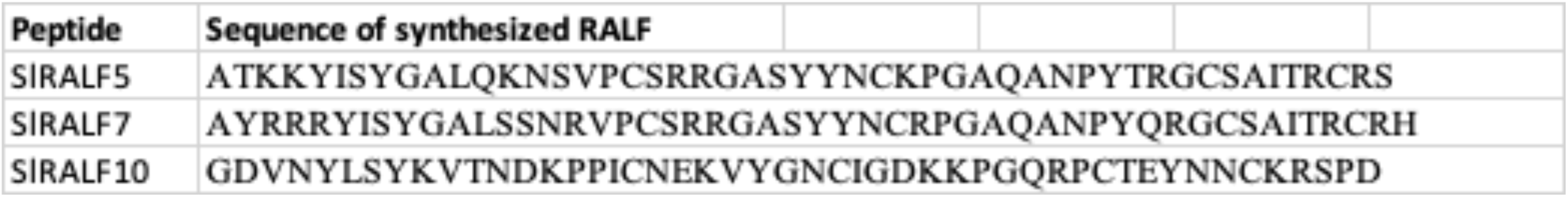
Amino acid sequence of the synthesized *Sl*RALF peptides.

| Peptide | Sequence of synthesized RALF |
| --- | --- |
| *SIRALF5* | ATKKYISYGALQKNSVPCSRRGASYYNCKPGAQANPYTRGCSAITRCRS |
| *SIRALF7* | AYRRRYISYGALSSNRVPCSRRGASYYNCRPGAQANPYQRGCSAITRCRH |
| *SIRALF10* | GDVNYLSYKVTNDKPPICNEKVYGNCIGDKKPGQRPCTEYNNCKRSPD |

**Table S4:**
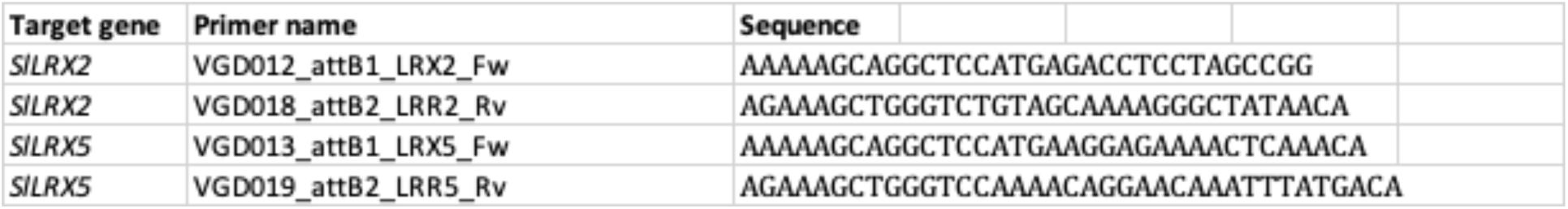
Cloning primers for tomato genomic DNA.

## References

Abarca, A., Franck, C. M., & Zipfel, C. (2021). Family-wide evaluation of rapid alkalinization factor peptides. Plant Physiology, 187(2), 996–1010. 10.1093/plphys/kiab308

Abramson, J., Adler, J., Dunger, J., Evans, R., Green, T., Pritzel, A., Ronneberger, O., Willmore, L., Ballard, A. J., Bambrick, J., Bodenstein, S. W., Evans, D. A., Hung, C. C., O’Neill, M., Reiman, D., Tunyasuvunakool, K., Wu, Z., Žemgulytė, A., Arvaniti, E., … Jumper, J. M. (2024). Accurate structure prediction of biomolecular interactions with AlphaFold 3. Nature, 630(8016), 493–500. 10.1038/s41586-024-07487-w

Amorim-Silva, V., García-Moreno, Á., Castillo, A. G., Lakhssassi, N., Del Valle, A. E., Pérez-Sancho, J., Li, Y., Posé, D., Pérez-Rodriguez, J., Lin, J., Valpuesta, V., Borsani, O., Zipfel, C., Macho, A. P., & Botella, M. A. (2019). TTL proteins scaffold brassinosteroid signaling components at the plasma membrane to optimize signal transduction in arabidopsis. Plant Cell, 31(8), 1807–1828. 10.1105/tpc.19.00150

Atkinson, R. G., Sutherland, P. W., Johnston, S. L., Gunaseelan, K., Hallett, I. C., Mitra, D., Brummell, D. A., Schröder, R., Johnston, J. W., & Schaffer, R. J. (2012). Down-regulation of POLYGALACTURONASE1 alters firmness, tensile strength and water loss in apple (Malus x domestica) fruit. BMC Plant Biology, 12. 10.1186/1471-2229-12-129

Bedinger, P. A., Pearce, G., & Covey, P. A. (2010). RALFs. Plant Signaling & Behavior, 5(11), 1342–1346. 10.4161/psb.5.11.12954

Blackburn, M. R., Haruta, M., & Moura, D. S. (2020). Twenty years of progress in physiological and biochemical investigation of RALF peptides. Plant Physiology 182 (4), 1657–1666. 10.1104/PP.19.01310

Brummell, D. A. (2006). Cell wall disassembly in ripening fruit. Functional Plant Biology 33 (2), 103–119. 10.1071/FP05234

Campbell, L., & Turner, S. R. (2017). A comprehensive analysis of RALF proteins in green plants suggests there are two distinct functional groups. Frontiers in Plant Science, 8. 10.3389/fpls.2017.00037

Cesarino, I. (2023). Killing me softly: A pathogen accelerates fruit ripening and softening to cause disease. Plant Physiology, 191(1), 21–23. 10.1093/plphys/kiac469

Ge, Z., Bergonci, T., Zhao, Y., Zou, Y., Du, S., Liu, M.-C., Luo, X., Ruan, H., García-Valencia, L. E., Zhong, S., Hou, S., Huang, Q., Lai, L., Moura, D. S., Gu, H., Dong, J., Wu, H.-M., Dresselhaus, T., Xiao, J., … Qu, -Jia. (2017). Arabidopsis pollen tube integrity and sperm release are regulated by RALF-mediated signaling. Science 358 (6370), 1596–1600. 10.1126/science.aao3642.

Herger, A., Dünser, K., Kleine-Vehn, J., & Ringli, C. (2019). Leucine-Rich Repeat Extensin Proteins and Their Role in Cell Wall Sensing. In Current Biology (Vol. 29, Number 17, pp. R851–R858). Cell Press. 10.1016/j.cub.2019.07.039

Huang, X., Liu, Y., Jia, Y., Ji, L., Luo, X., Tian, S., & Chen, T. (2024). FERONIA homologs in stress responses of horticultural plants: current knowledge and missing links. Stress Biology 4 (1). 10.1007/s44154-024-00161-1

Hyodo, H., Terao, A., Furukawa, J., Sakamoto, N., Yurimoto, H., Satoh, S., & Iwai, H. (2013). Tissue specific localization of pectin-Ca2+ cross-linkages and pectin methyl-esterification during fruit ripening in tomato (Solanum lycopersicum). PLoS ONE, 8(11). 10.1371/journal.pone.0078949

Jeong, H. Y., Nguyen, H. P., Eom, S. H., & Lee, C. (2018). Integrative analysis of pectin methylesterase (PME) and PME inhibitors in tomato (Solanum lycopersicum): Identification, tissue-specific expression, and biochemical characterization. Plant Physiology and Biochemistry, 132, 557–565. 10.1016/j.plaphy.2018.10.006

Ji, D., Cui, X., Qin, G., Chen, T., & Tian, S. (2020). SlFERL interacts with S-adenosylmethionine synthetase to regulate fruit ripening. Plant Physiology, 184(4), 2168–2181. 10.1104/pp.20.01203

Ji, D., Liu, W., Cui, X., Liu, K., Liu, Y., Huang, X., Li, B., Qin, G., Chen, T., & Tian, S. (2023). A receptor-like kinase SlFERL mediates immune responses of tomato to Botrytis cinerea by recognizing BcPG1 and fine-tuning MAPK signaling. New Phytologist, 240(3), 1189–1201. 10.1111/nph.19210

Lee, H. K., & Santiago, J. (2023). Structural insights of cell wall integrity signaling during development and immunity. Current Opinion in Plant Biology, 76. 10.1016/j.pbi.2023.102455

Li, R., Wang, X., Haj Ahmad, F., Fuglsang, A. T., Steppuhn, A., Stintzi, A., & Schaller, A. (2025). Poltergeist-Like 2 (PLL2)-dependent activation of herbivore defence distinguishes systemin from other immune signalling pathways. In Nature Plants (Vol. 11, Number 7, pp. 1270–1281). Nature Research. 10.1038/s41477-025-02040-7

Liu, L., Liu, X., Bai, Z., Tanveer, M., Zhang, Y., Chen, W., Shabala, S., & Huang, L. (2024). Small but powerful: RALF peptides in plant adaptive and developmental responses. Plant Science, 343. 10.1016/j.plantsci.2024.112085

Lu, R., Lanooij, J., & Smakowska-Luzan, E. (2025). From roots to reproduction: the multifaceted roles of rapid alkalinization factor and epidermal patterning factor peptides in plants. Journal of Experimental Botany 76 (19) 5713–5727. 10.1093/jxb/eraf303

Ma, W., Liu, X., Chen, K., Yu, X., & Ji, D. (2023). Genome-Wide Re-Identification and Analysis of CrRLK1Ls in Tomato. International Journal of Molecular Sciences, 24(4). 10.3390/ijms24043142

Manchanda, P., & Geitmann, A. (2025). The tri-molecular interaction controlling plant cell structure. Current Opinion in Cell Biology, 95. 10.1016/j.ceb.2025.102538

Mecchia, M. A., Santos-Fernandez, G., Duss, N. N., Somoza, S. C., Boisson-Dernier, A., Gagliardini, V., Martínez-Bernardini, A., Fabrice, T. N., Ringli, C., Muschietti, J. P., & Grossniklaus, U. (2017). RALF4/19 peptides interact with LRX proteins to control pollen tube growth in Arabidopsis. Science 358 (6370); 1600–1603. 10.1126/science.aao5467.

Merino, M. C., Guidarelli, M., Negrini, F., De Biase, D., Pession, A., & Baraldi, E. (2019). Induced expression of the Fragaria × ananassa Rapid alkalinization factor-33-like gene decreases anthracnose ontogenic resistance of unripe strawberry fruit stages. Molecular Plant Pathology, 20(9), 1252–1263. 10.1111/mpp.12837

Montano, J. A., Posé, S., Mercado, J. A., & Doblas, V. G. (2026). Insights into the roles of RALF peptides and *Cr*RLK1L receptors in fruit ripening. The Plant Journal, 125(5). 10.1111/tpj.70766

Morato do Canto, A., Ceciliato, P. H. O., Ribeiro, B., Ortiz Morea, F. A., Franco Garcia, A. A., Silva-Filho, M. C., & Moura, D. S. (2014). Biological activity of nine recombinant AtRALF peptides: Implications for their perception and function in Arabidopsis. Plant Physiology and Biochemistry, 75, 45–54. 10.1016/j.plaphy.2013.12.005

Moussu, S., Broyart, C., Santos-Fernandez, G., Augustin, S., Wehrle, S., Grossniklaus, U., & Santiago, J. (2020). Structural basis for recognition of RALF peptides by LRX proteins during pollen tube growth. Proceedings of the National Academy of Sciences of the United States of America, 117(13), 7494–7503. 10.1073/pnas.2000100117

Moussu, S., & Ingram, G. (2023). The EXTENSIN enigma. The Cell Surface, 9. 10.1016/j.tcsw.2023.100094

Moussu, S., Lee, H. K., Haas, K. T., Broyart, C., Rathgeb, U., De Bellis, D., Levasseur, T., Schoenaers, S., Fernandez, G. S., Grossniklaus, U., Bonnin, E., Hosy, E., Vissenberg, K., Geldner, N., Cathala, B., Höfte, H., & Santiago, J. (2023). Plant cell wall patterning and expansion mediated by protein-peptide-polysaccharide interaction. Science, 382(6671), 719–725. 10.1126/science.adi4720

Murphy, E., & De Smet, I. (2014). Understanding the RALF family: A tale of many species. Trends in Plant Science, 19 (10), 664–671. 10.1016/j.tplants.2014.06.005

Negrini, F., O’Grady, K., Hyvönen, M., Folta, K. M., & Baraldi, E. (2020). Genomic structure and transcript analysis of the Rapid Alkalinization Factor (RALF) gene family during host-pathogen crosstalk in Fragaria vesca and Fragaria x ananassa strawberry. PLoS ONE, 15(3). 10.1371/journal.pone.0226448

Palin, R., & Geitmann, A. (2012). The role of pectin in plant morphogenesis. BioSystems, 109(3), 397–402. 10.1016/j.biosystems.2012.04.006

Paniagua, C., Blanco-Portales, R., Barceló-Muñoz, M., García-Gago, J. A., Waldron, K. W., Quesada, M. A., Muñoz-Blanco, J., & Mercado, J. A. (2016). Antisense down-regulation of the strawberry β-galactosidase gene FaβGal4 increases cell wall galactose levels and reduces fruit softening. Journal of Experimental Botany, 67(3), 619–631. 10.1093/jxb/erv462

Pearce, G., Yamaguchi, Y., Munske, G., & Ryan, C. A. (2010). Structure-activity studies of RALF, Rapid Alkalinization Factor, reveal an essential - YISY - motif. Peptides, 31(11), 1973–1977. 10.1016/j.peptides.2010.08.012

Peng, Z., Liu, G., Li, H., Wang, Y., Gao, H., Jemrić, T., & Fu, D. (2022). Molecular and Genetic Events Determining the Softening of Fleshy Fruits: A Comprehensive Review. International Journal of Molecular Sciences, 23 (20). 10.3390/ijms232012482

Posé, S., Paniagua, C., Matas, A. J., Gunning, A. P., Morris, V. J., Quesada, M. A., & Mercado, J. A. (2019). A nanostructural view of the cell wall disassembly process during fruit ripening and postharvest storage by atomic force microscopy. Trends in Food Science and Technology, 87, 47–58. 10.1016/j.tifs.2018.02.011

Pratyusha, D. S., & Sarada, D. V. L. (2025). Rapid Alkalinization Factor – A cryptide regulating developmental and stress responses. Plant Science, 359. 10.1016/j.plantsci.2025.112600

Qian, M., Xu, Z., Zhang, Z., Li, Q., Yan, X., Liu, H., Han, M., Li, F., Zheng, J., Zhang, D., & Zhao, C. (2021). The downregulation of PpPG21 and PpPG22 influences peach fruit texture and softening. Planta, 254(2). 10.1007/s00425-021-03673-6

Sang, Y. & Macho, A.P. (2017). Analysis of PAMP-triggered ROS burst in plant immunity. Methods Molecular Biology, 1578, 143–153. 10.1007/978-1-4939-6859-6_11.

Santiago-Doménech, N., Jiménez-Bemúdez, S., Matas, A. J., Rose, J. K. C., Muñoz-Blanco, J., Mercado, J. A., & Quesada, M. A. (2008). Antisense inhibition of a pectate lyase gene supports a role for pectin depolymerization in strawberry fruit softening. Journal of Experimental Botany, 59(10), 2769–2779. 10.1093/jxb/ern142

Sato, S., Tabata, S., Hirakawa, H., Asamizu, E., Shirasawa, K., Isobe, S., Kaneko, T., Nakamura, Y., Shibata, D., Aoki, K., Egholm, M., Knight, J., Bogden, R., Li, C., Shuang, Y., Xu, X., Pan, S., Cheng, S., Liu, X., … Gianese, G. (2012). The tomato genome sequence provides insights into fleshy fruit evolution. Nature, 485(7400), 635–641. 10.1038/nature11119

Schade, S., Leicher, H., von Arx, M., Monte, I., Gronnier, J., & Stegmann, M. (2025). The interplay of RALF structural and signaling functions in plant-microbe interactions. PLOS Pathogens, 21(10). 10.1371/journal.ppat.1013588

Schoenaers, S., Lee, H. K., Gonneau, M., Faucher, E., Levasseur, T., Akary, E., Claeijs, N., Moussu, S., Broyart, C., Balcerowicz, D., AbdElgawad, H., Bassi, A., Damineli, D. S. C., Costa, A., Feijó, J. A., Moreau, C., Bonnin, E., Cathala, B., Santiago, J., … Vissenberg, K. (2024). Rapid alkalinization factor 22 has a structural and signalling role in root hair cell wall assembly. Nature Plants, 10(3), 494–511. 10.1038/s41477-024-01637-8

Sénéchal, F., Wattier, C., Rustérucci, C., & Pelloux, J. (2014). Homogalacturonan-modifying enzymes: Structure, expression, and roles in plants. Journal of Experimental Botany, 65(18), 5125–5160. 10.1093/jxb/eru272

Shi, Y., Li, B. J., Grierson, D., & Chen, K. S. (2023). Insights into cell wall changes during fruit softening from transgenic and naturally occurring mutants. Plant Physiology, 192(3), 1671–1683. 10.1093/plphys/kiad128

Solis-Miranda, J., & Quinto, C. (2021). The CrRLK1L subfamily: One of the keys to versatility in plants. Plant Physiology and Biochemistry 166, 88–102. 10.1016/j.plaphy.2021.05.028

Srivastava, R., Liu, J. X., Guo, H., Yin, Y., & Howell, S. H. (2009). Regulation and processing of a plant peptide hormone, AtRALF23, in Arabidopsis. Plant Journal, 59(6), 930–939. 10.1111/j.1365-313X.2009.03926.x

Uluisik, S., & Seymour, G. B. (2020). Pectate lyases: Their role in plants and importance in fruit ripening. Food Chemistry, 309. 10.1016/j.foodchem.2019.125559

Wang, D., & Seymour, G. B. (2022). Molecular and biochemical basis of softening in tomato. Molecular Horticulture, 2 (1). 10.1186/s43897-022-00026-z

Wang, D., Yeats, T. H., Uluisik, S., Rose, J. K. C., & Seymour, G. B. (2018). Fruit Softening: Revisiting the Role of Pectin. Trends in Plant Science, 23 (4), 302–310. 10.1016/j.tplants.2018.01.006

Wang, X., Li, R., Stintzi, A. Schaller A. (2024). Automated real-time monitoring of extracellular pH to asses early plant defense signaling. Methods Molecular Biology, 2731, 157–167. 10.1007/978-1-0716-3511-7_12

Wang, P., He, T., Xiao, S., Gao, Y., & Li, Z. (2025). Genome-wide identification and characterization of Rapid alkalinization factors (RALFs) in tomato and function analysis of SlRALF2/3 in immunity. Plant Stress, 18. 10.1016/j.stress.2025.101090

Xiao, Y., Stegmann, M., Han, Z., DeFalco, T. A., Parys, K., Xu, L., Belkhadir, Y., Zipfel, C., & Chai, J. (2019). Mechanisms of RALF peptide perception by a heterotypic receptor complex. Nature, 572(7768), 270–274. 10.1038/s41586-019-1409-7

Zhang, H., Jing, X., Chen, Y., Liu, Z., Xin, Y., & Qiao, Y. (2020). The genome-wide analysis of RALF-like genes in strawberry (Wild and cultivated) and five other plant species (rosaceae). Genes, 11(2). 10.3390/genes11020174

Zhang, R., Shi, P. T., Zhou, M., Liu, H. Z., Xu, X. J., Liu, W. T., & Chen, K. M. (2023). Rapid alkalinization factor: function, regulation, and potential applications in agriculture. Stress Biology, 3, (1). 10.1007/s44154-023-00093-2

Zhu, S., Fu, Q., Xu, F., Zheng, H., & Yu, F. (2021). New paradigms in cell adaptation: decades of discoveries on the CrRLK1L receptor kinase signalling network. New Phytologist 232 (3), 1168–1183. 10.1111/nph.17683

